# Blue spectral quality contributes to yield and ascorbic acid content in ‘Micro Tom’ tomato under sole-source lighting

**DOI:** 10.1101/2023.11.27.568869

**Authors:** Matthew S. Goldman, Elsebeth Kolmos

## Abstract

There is an intricate relationship between the spectral composition of light, plant photosynthetic performance and biomass accumulation. The interaction between plants and the ambient light environment is not only crop-specific but also crucial for maximized yield and nutritional quality. The emergence of LED technology introduced a unique avenue for manipulating the light spectrum, offering new and continued possibilities for optimizing growth conditions. With a focus on tomato cultivation under sole-source lighting, we have investigated the impact of blue spectral quality (waveband composition) on growth and nutritional value of dwarf tomato. Notably, our study revealed that distinct wavebands of blue light (400 nm, 420 nm and 450 nm) can influence the net photosynthetic rate despite unaltered photochemical efficiency. Our research uncovered a correlation whereby shorter wavelengths of blue light increased leaf area, while longer blue wavelengths contributed to greater harvest indices. In addition, we identified a specific blue peak wavelength, 450 nm, that significantly affected chlorophyll composition in leaves and ascorbate levels in fruits. Through these findings, we call attention to the notion that blue spectral quality has a role in shaping both yield and nutritional attributes of dwarf tomato, such as ‘Micro Tom’, under sole-source lighting. Overall, our research provides valuable insight into the nuanced interplay between light spectrum, plant physiology, and horticultural outcome.

## 1. Introduction

Tomato is a diverse crop that can be grown in the field or indoors in a controlled environment. Due to the excellent nutritional and organoleptic properties, the tomato fruit is very popular, both as a fresh and a processed food. Therefore, tomato has recently been the most abundantly produced fruit worldwide (2.57·10^8^ t in 2021, compared to ∼1.38·10^8^ t each for apples and bananas; FAOSTAT, 2021).

Recently, the focus of the indoor or vertical farming industry has been to move beyond the production of leafy greens. Expanding the crop production in indoor vertical farms to include fruits, non-leafy vegetables and value-added crops would enhance the industry’s economic sustainability by offering a more diverse range of products for consumers, i.e., also benefiting nutrition and food security (Hikosaka, 2018; Zelkind et al., 2022).

Tomato fruits produced in the greenhouse and other indoor operations are mainly sold for fresh consumption. The important nutritional components of tomato are secondary metabolites such as carotenoids, vitamin C (ascorbic acid, ASA) and phenolic acids, which all act as antioxidants. Tomato is a key source of the carotenoid lycopene, which gives the fruit its characteristically red color and is a popular antioxidant with anti-cancer properties (Salehi et al., 2019). ASA is an essential nutrient for humans. Tomato, therefore, despite its relatively low ascorbate content compared to oranges and grapes, is a key food – so widely consumed that it has become the number three of fruits important for ASA intake (Fenech et al., 2018).

Crop production in indoor or vertical farms offers the opportunity to fully shape plant growth due to complete environmental control (SharathKumar et al., 2020). In this regard, the quantity, quality and periodicity of light is important (de Carbonnel et al., 2022). In vertical farms, sole source lighting, such as light emitting diodes (LED), is substituted for sunlight (Oh and Lu, 2022). LED lights come in many colors, from UV (∼300-400 nm) to far-red (∼750 nm) light, and can save energy cost compared to other lamp types, e.g., fluorescent and high-pressure sodium (HPS) lights. The energy efficiency, ratio of electrical input to photon output energy, of the individual LEDs differ, for example the blue LEDs are the most efficient (49% vs. 32% for red LEDs) (Mitchell, 2022). Examples of simple LED grow lights are fixtures populated entirely with white broadband or only blue and red LED chips. For versatility, albeit more costly, the fixture can consist of a mix of broad- and narrowband LEDs to enable customizable spectra (Nozue and Gomi, 2018).

For grow-light recipes, using single color LEDs, the basic design is blue (400-500 nm) combined with red (600-700 nm) light (resulting in magenta) because blue and red are the main wavelength bands in the photosynthetically active radiation spectrum (PAR, 400-700 nm) and crucial to proper plant growth and development. Specifically, blue light is important for elongation growth and red light for high quantum efficiency of photosynthesis. In addition, inclusion of green (500-600 nm) and far-red (700-800 nm) light is desirable for increased photosynthetic efficiency (Paradiso and Proietti, 2021). Far-red light is included in the extended PAR (ePAR, 400-750 nm) spectrum and is also important for leaf expansion (Zhen et al., 2021). In addition to plant morphology, light quality also affects the content of secondary metabolites in plants including fruit quality, which reflects crop nutritional value and consumer appeal (Ouzounis et al., 2015).

Blue light spans the wavelengths of 400-500 nm. 400 nm borders the UV-A (320-400 nm) spectrum, which can promote plant stress responses including non-green pigment formation, however this is mostly known for shorter UV wavelengths than UV-A radiation (Artés-Hernández et al., 2022). Blue light regulates plant growth (cell elongation) and promotes photosynthetic efficiency and stomatal conductance (Cosgrove, 1981; Matsuda et al., 2004). Without blue light leaves and stems are elongated, reminiscent of the shade-avoidance response and termed “red light syndrome”, and photosystem synthesis is impaired (Mochizuki et al., 2004; Muneer et al., 2014). Such developmental defects can be prevented with a blue light amount of 7-8% of the photon flux density (PFD) (Hogewoning et al., 2010; Kaiser et al., 2018). Tomato-specific blue light benefits include prevention of grey mold disease and reduction of intumescence injury (Imada et al., 2014; Retana-Cordero et al., 2022). In addition, blue light affects tomato nutritional quality (He et al., 2022; Ntagkas et al., 2020; Vitale et al., 2022; Wang et al., 2022).

Since long-term analysis of spectral quality at the whole plant level is rare (Sytar et al., 2021), including for tomato, e.g., limited to studies by Kalaitzoglou et al. (2021), Ke et al. (2022 and 2023) and Utasi et al. (2023), we here sought to compare the effect of different blue LED wavebands (with peaks at 400 nm, 420 nm and 450 nm) on the growth and yield of tomato under sole-source lighting. Our study had two major objectives. The first one was a comparative analysis of photosynthetic CO_2_ assimilation under mono- and poly-chromatic light conditions following changes in the blue spectral wavebands. The second was analysis of growth and yield of tomato under poly-chromatic light following similar changes in the blue part of the spectrum as for part one of our study. We found that only the net photosynthetic rate, not the photochemical efficiency, responded to blue light changes when measured in isolation at the single leaf level. The growth and yield of tomato was even more responsive to the blue spectral quality than photosynthesis. Short-wavelength blue light promoted leaf size whereas longer blue wavelengths resulted in increased biomass and fruit/leaf ratio (harvest index). Specifically, blue light with a peak wavelength of 450 nm was associated with relatively low leaf chlorophyll (Chl) a/b ratio and high ASA content in the tomato fruits. Our study revealed that blue spectral quality has a subtle influence on leaf photochemistry compared to the effect on tomato crop physiological outputs.

## 2. Materials and methods

### 2.1. Plant material, cultivation and harvest

We used the dwarf tomato (*Solanum lycopersicum*) ‘Micro Tom’, which is a tomato model plant (Scott, 1989). The seeds were purchased from Tomato Growers Supply Company (Fort Myers, FL). Each growth trial had twelve plants that were reduced to four plants after one month of growth. The growth trials were randomized between five indoor growth cabinets (A1000, Conviron, Winnipeg, Canada) and each trial was repeated three times. The seeds were germinated on wet substrate (Rockwool® cubes) and transferred to the trial conditions at the 3-leaf stage. The plants were cultivated in hydroponic deep-water reservoirs with half strength modified Hoagland’s solution (Hoagland and Arnon, 1950) containing the following fertilizer elements: KOH (0.8 mM), KNO_3_ (4 mM), Ca(NO_3_)_2_ (14 mM), MgSO_4_ (7.5 mM), NH_4_PO_4_ (2 mM), FeSO_4_·7H_2_O (1 mM), H_3_BO_3_ (9 μM), MnCl_2_·4 H_2_O (10 μM), ZnSO_4_·7H_2_O (8 μM), CuSO_4_·5H_2_O (6 μM), and (NH_4_)_6_Mo_7_O_24_·4H_2_O (0.08 mM). The reservoirs had aeration and the fertilizer solution was refreshed as required to maintain electrical conductivity of 1.2 dS m^-1^. CO_2_ levels were ambient and the relative humidity was ∼60%. The diurnal temperature regime was 25°C/18°C. When two thirds of the fruit were ripe, at 120 days after sowing (DAS), the plants were harvested and cut at the base of the stem. The organs were separated, measured and weighed using an analytical balance (Mettler Toledo) before and after oven drying (Heratherm, ThermoFisher Scientific, Waltham, MA) for 2-3 days (until no change in weight). The roots were not analyzed. Leaf area was determined using a laser area meter (CI-202, CID Bio-Science). All fruits were counted and measured, and the red fruits and leaves were stored at - 80°C until further chemical analysis. The dry-weight of the fruit was not determined. Unripe (non-red) fruit was not chemically analyzed.

### 2.2. Light treatment

The poly-chromatic LED spectrum, used for light response curves and subsequent growth trials (applied after the seedling stage), was 9% blue light (with peak wavelengths of 400 nm, 420 nm or 450 nm; Figure 1), 18% green (with a peak wavelength of 530 nm), 64% red (with a peak wavelength of 660 nm) and 9% far-red (with a peak wavelength of 735 nm) light using TIGER lights with individual dimmable LED channels (Karlicek, 2019). The intensity was 400 μmol m^-2^ s^-1^ at the base of the plants and was not adjusted during growth. The photoperiod was 16 hours and the daily light integral (DLI) 23 mol m^-2^ d^-1^. The light settings were confirmed with a spectrometer (JAZ, OceanOptics, Rochester, NY) and ePAR sensor (Apogee Instruments, Logan, UT).

**Figure 1.**
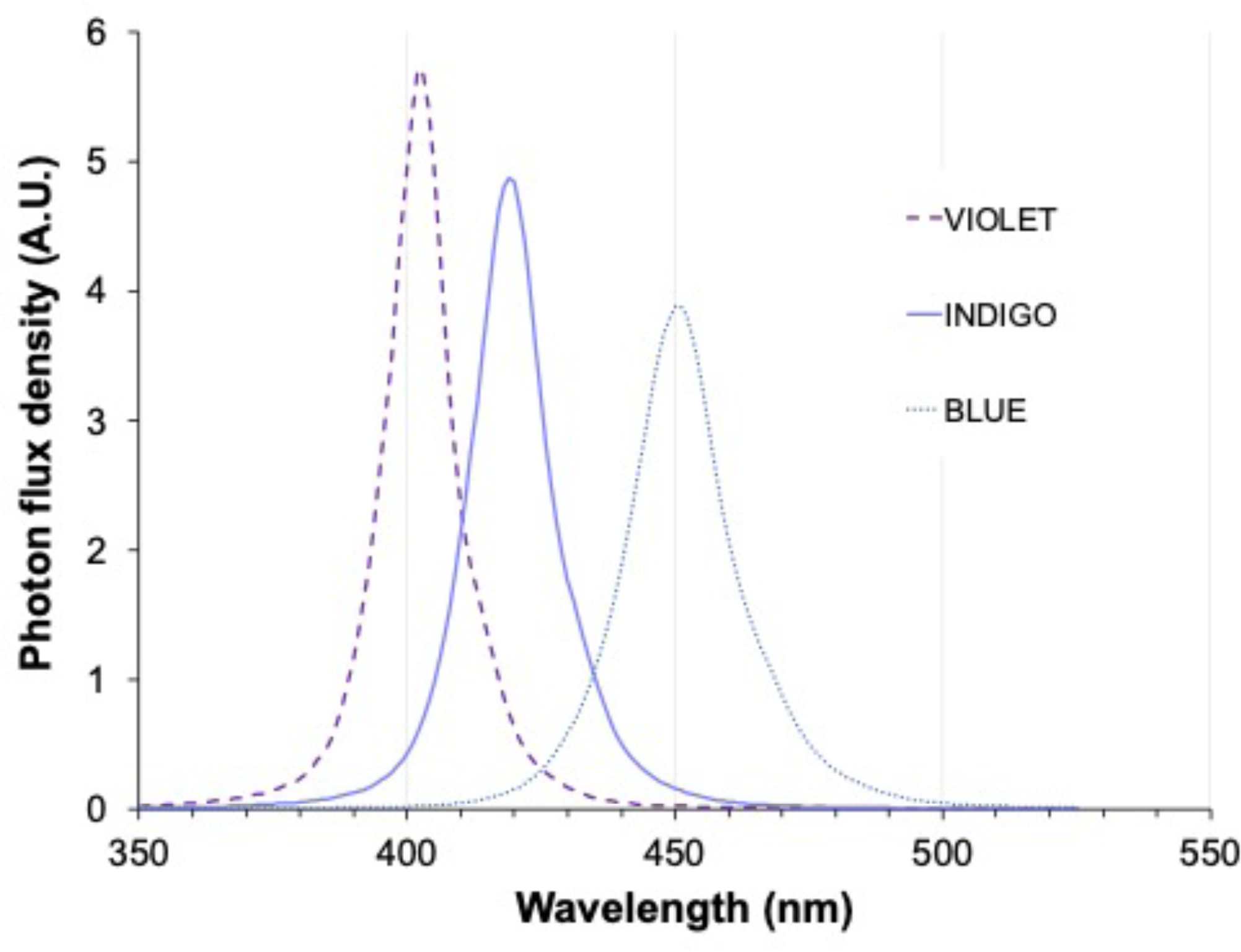
The three blue LED spectra evaluated. The wavebands included VIOLET (with a peak wavelength of 400 nm, FWHM = 12 nm), INDIGO (with a peak wavelength of 420 nm, FWHM = 15 nm) and BLUE (with a peak wavelength of 450 nm, FWHM = 15 nm) of the TIGER LED light. Note, both VIOLET and INDIGO extends outside the PAR spectrum, which is 400-700 nm.

### 2.3. Gas exchange measurement

For light response curves, the gas exchange rate was measured in response to step decreases in light intensities of the spectrum- or wavelength-of-interest with the infra-red gas analyzer (IRGA; CIRAS-3, PP Systems, Amesbury, MA) equipped with the TIGER Cub LED attachment (Kolmos et al., 2021). During readings, the ambient CO_2_ concentration was kept steady at ∼400 μmol μmol^-1^ (by the IRGA instrument) and the ambient temperature was 22°C inside the leaf cuvette. For each light intensity, leaf gas exchange from 3-5 plants was measured, using the youngest fully expanded leaf. The analysis of the light response (P_N_/I) curves was used for the estimation of the maximum light saturated photosynthetic rate (P_gmax_), apparent quantum efficiency (ϕ), light compensation (I_comp_), light saturation point (I_sat_), and the dark respiration R_D_. Curve fitting was performed with the least squares method using the Excel tool where the nine tool models were tested for the best fit to the data (Lobo et al., 2013). Plants were 3-4 weeks old and grown under fluorescent light conditions (Philips T5) prior to the photosynthetic measurements. During growth trials, IRGA readings were performed on day 40.

### 2.4. Chlorophyll a fluorescence measurement

The chlorophyll *a* fluorescence (ChlF) readings were performed with a portable chlorophyll fluorometer (PAM-2500, Walz, Effeltrich, Germany) on the most recent fully developed leaf. Plants were dark-adapted for 20 min and then illuminated with a saturating pulse of the default red light of the PAM. After 1 min red, actinic light was turned on for 15-20 min. Saturating pulses of 125 μmol m^-2^ s^-1^ were applied to measure quenched fluorescence values from the leaf. The ChlF measurement was conducted on 10 plants from each light treatment.

### 2.5. Leaf pigment analysis

The chlorophyll and total carotenoid content (TCC) from methanolic leaf extracts of pooled frozen leaf tissue was analyzed using UV-VIS spectrophotometry (V-570, Jasco, Easton, MD) according to Lichtenthaler and Buschmann (2001).

### 2.6. Fruit nutritional analysis

All nutritional assays were performed using homogenized fruit. Lycopene and beta-carotene content was analyzed simultaneously by UV-VIS spectrophotometry from fruit extracts using 4:6 (vol:vol) acetone-hexane solvent according to the protocol of Nagata and Yamashita (1992). The ASA content of tomato fruits was determined using a titration assay and standard curve with commercial ascorbic acid (S88476, Sigma) and 2,6-dichloroindophenol (119814, Sigma) (AOAC, 2006). Total phenol content (TPC) was determined with reference to a standard curve with gallic acid and using Folin-Ciocalteu reagent (F9252, Sigma) added to methanolic fruit extracts, according to the protocol of Ainsworth and Gillespie (2007). The measure of total soluble solids (TSS; °Brix) was determined using a handheld refractometer (PAL-1, Atago, Bellevue, WA).

### 2.7. Data analysis

Based on independent biological triplicates (trials), each with 4-10 technical replicates (plants), experimental results were expressed as the mean with standard error of the mean (S.E.M.) and graphed in Prism 10 software (GraphPad, San Diego, CA). Also, using this software, the data were analyzed using nested one-way ANOVA and the different means were assessed using Tukey’s multiple comparisons test with significance level as indicated.

## 3. Results

### 3.1. Light response curves

To characterize the photosynthetic response of tomato with relation to changes in blue spectral quality, we performed single leaf infra-red gas exchange measurements over a range of light intensities (from high to low fluence rate). We compared the resulting light response of CO_2_ assimilation rate for the three blue wavelength bands, VIOLET (with a peak wavelength of 400 nm), INDIGO (with a peak wavelength of 420 nm) and BLUE (with a peak wavelength of 450 nm), under mono- and poly-chromatic conditions (Figure 2A and B).

**Figure 2.**
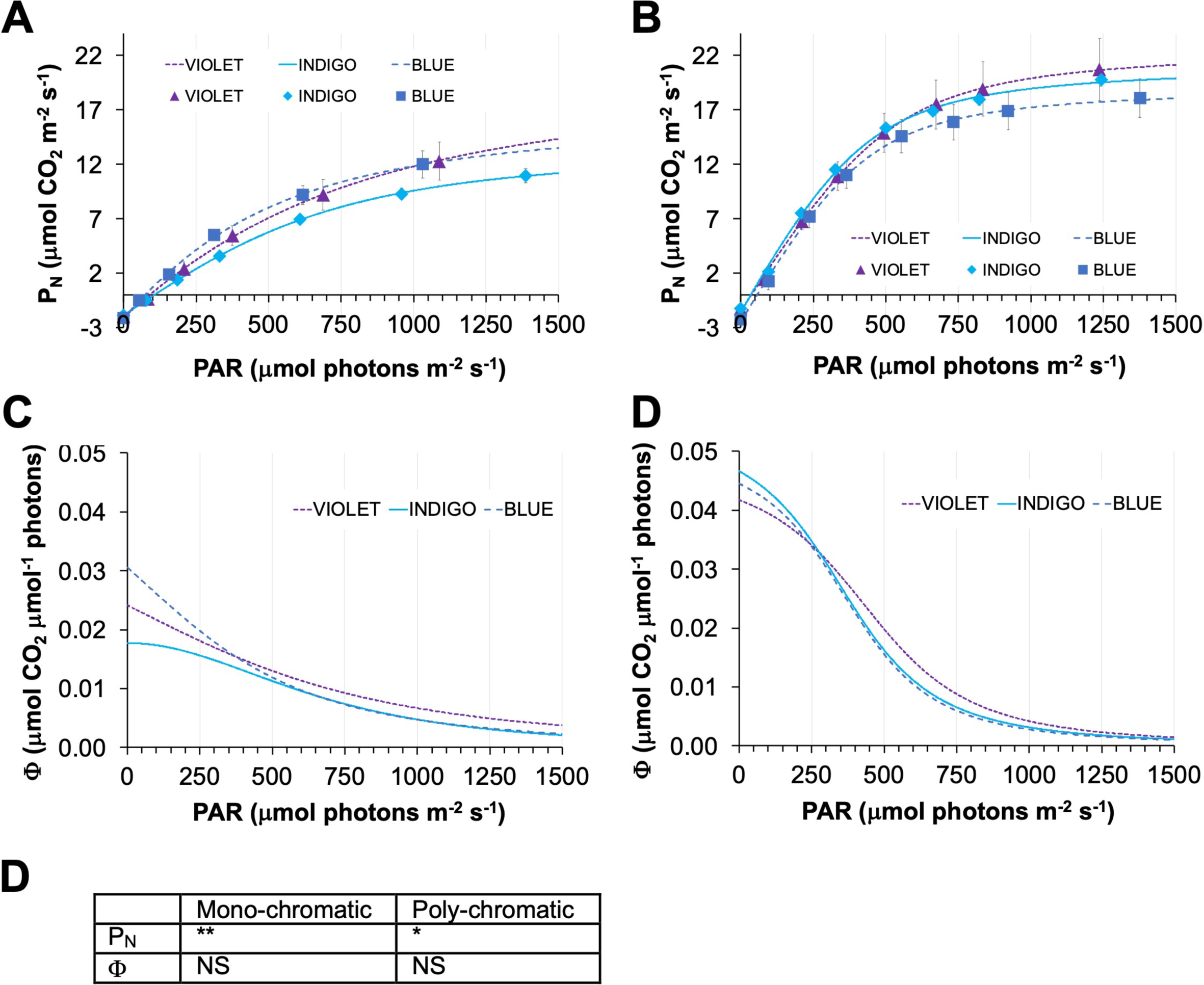
Light response curve analysis. Tomato single leaf light response for blue light with wavelengths 400 nm (VIOLET), 420 nm (INDIGO) and 450 nm (BLUE). (A) Light response curves of the net photosynthetic rate, P_N_, for mono- and (B) poly-chromatic light. (C) The estimated quantum efficiency, Φ, light response for mono- and (D) poly-chromatic light. The solid lines represent fitted curves based on the average of 3 independent biological measurements. Error bars are standard error of the mean, S.E.M. (E) Repeated measures ANOVA results for the measured (P_N_) and estimated (Φ) variables of tomato light response curves. P_N_, net photosynthetic rate (P_gross_ – R_D_); <λ, photochemical efficiency. *, *P* < 0.05; **, *P* < 0.01.

We found that changing the blue wavebands in both mono- and poly-chromatic spectra affected different parameters of the tomato light response curves (Figure 2, Table 1 and 2). The net photosynthetic rate, P_N_, but not the corresponding estimated quantum efficiency (Phi, Φ) of PSII, was significantly different within mono- and poly-chromatic spectra (Figure 2 and Table 1). Thus, even though the blue spectrum did not appear to influence PSII photochemical performance, the rate of net photosynthesis was affected.

**Table 1.** Tomato photosynthetic light response data. The values are based on the fitted regression curves in Figure 2, calculated according to (Lobo et al., 2013). CV, coefficient of variance; R_D_, dark respiration, μmol CO_2_ m^-2^ s^-1^; I, light intensity, μmol m^-2^ s^-1^; I_max_, light saturation point beyond which there is no significant change in P_N_, μmol m^-2^ s^-1^; P_N_, net photosynthetic rate at maximum light intensity, μmol CO_2_ m^-2^ s^-1^; LCP, light compensation point at I = 0, μmol photons m^-2^ s^-1^; LSP_90%_, light saturation point at 90% of I_max_, μmol photons m^-2^ s^-1^; <λ_LCP_, photochemical efficiency at LCP, μmol CO_2_ μmol^-1^ photons; <λ_400_, photochemical efficiency at I_400_, μmol CO_2_
μmol^-1^ photons.

|  | Mono-chromatic spectrum (with peaks at) |  |  |  |  | Poly-chromatic spectrum (with blue peaks at) |  |  |  |  |
| --- | --- | --- | --- | --- | --- | --- | --- | --- | --- | --- |
|  | VIOLET (400 nm) | INDIGO (420 nm) | BLUE (450 nm) | Mean value | CV (%) | VIOLET (400 nm) | INDIGO (420 nm) | BLUE (450 nm) | Mean value | CV (%) |
| $R_D$ | 2.04 | 1.84 | 2.12 | 2.00 | 7.21 | 1.48 | 1.56 | 2.44 | 1.83 | 29.2 |
| $P_N (I_{\text{max}})$ | 6.23 | 3.75 | 6.82 | 5.60 | 29.1 | 16.53 | 15.49 | 13.84 | 15.29 | 8.87 |
| LCP | 88.71 | 104.57 | 73.22 | 88.83 | 17.6 | 35.74 | 33.73 | 55.77 | 41.75 | 29.2 |
| $\text{LSP}_{90\%}$ | 5070.1 | 1879.1 | 3270.4 | 3406.5 | 47.0 | 1182.6 | 1064.6 | 1027.4 | 1091.5 | 7.42 |
| $\Phi_{\text{LCP}}$ | 0.0220 | 0.0173 | 0.0273 | 0.0222 | 22.5 | 0.0410 | 0.0456 | 0.0430 | 0.0432 | 5.34 |
| $\Phi_{400}$ | 0.0149 | 0.0131 | 0.0146 | 0.0142 | 6.79 | 0.0259 | 0.0232 | 0.0226 | 0.0239 | 7.36 |

For the mono-chromatic conditions, the dark respiration, R_D_, was very similar between all wavebands. The light compensation point (LCP) and the associated quantum efficiency (Φ_LCP_) was variable between the three blue wavebands but the variability was less than for P_N_(I_max_) and the light saturation point, LSP (Table 1).

For the poly-chromatic spectra, in contrast to the mono-chromatic conditions, most of the variation was seen for R_D_ and LCP. The other parameters, P_N_(I_max_), LSP and Φ, varied little between the different blue wavebands (Table 1).

The estimated quantum efficiency of PSII at 400 μmol m^-2^ s^-1^ (Φ_400_) had similar low variability (coefficient of variation, CV) among the three blue wavebands in mono- and polychromatic spectra (Table 1), indicating little effect of the blue spectral quality on quantum efficiency at the relevant irradiance for growing tomato plants.

### 3.2. Plant growth and yield

Based on the light response results (changes in P_N_), we sought to test the hypothesis that blue spectral quality is critical to tomato crop performance and yield, because these outputs are linked to photosynthetic performance (Long et al., 2006). For this we compared tomato growth and yield for the same poly-chromatic spectra used in the light response analysis.

During growth, chlorophyll *a* fluorescence (ChlF), CO_2_ and H_2_O exchange rates and morphological measurements were performed. ChlF measurements, including the PSII maximum efficiency (F_v_/F_m_), PSII quantum yield (Φ_PSII_), non-photochemical (NPQ) and photochemical quenching (qP), were similar for all spectra (Table 2). The gas exchanges rates were also largely unaltered, except for a slight P_N_ increase for BLUE compared to VIOLET and INDIGO (P<0.1; Table 3). In addition, no difference in plant height, stem diameter and chlorophyll content index was detected at bi-weekly measurements during growth (data not shown).

**Table 2.**
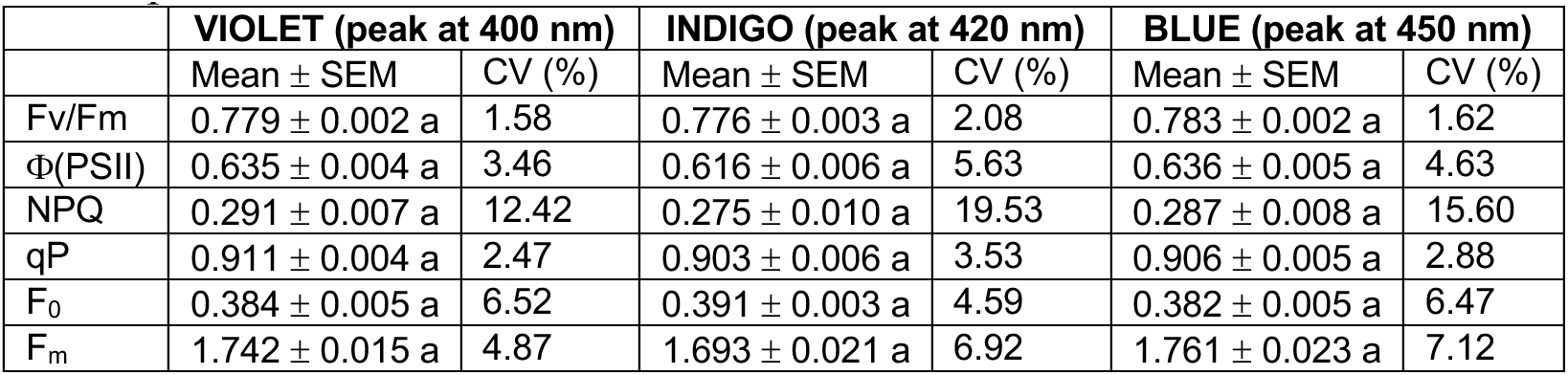
Tomato leaf ChlF measurement on day 40 during growth under poly-chromatic spectrum. Fv/Fm, maximum potential quantum efficiency of PSII photochemistry; Y(II), effective efficiency of PSII photochemistry; NPQ, non-photochemical quenching; qP, coefficient for photochemical efficiency; F_0_, minimum fluorescence; F_m_, maximum fluorescence. CV, coefficient of variance; SEM, standard error of the mean. The statistical significance of differences is indicated with letters (repeated measures one-way ANOVA, P<0.1). n = 10 for each experiment.

|  | VIOLET (peak at 400 nm) |  | INDIGO (peak at 420 nm) |  | BLUE (peak at 450 nm) |  |
| --- | --- | --- | --- | --- | --- | --- |
| | Mean ± SEM | CV (%) | Mean ± SEM | CV (%) | Mean ± SEM | CV (%) |
| $F_v/F_m$ | $0.779 \pm 0.002$ a | 1.58 | $0.776 \pm 0.003$ a | 2.08 | $0.783 \pm 0.002$ a | 1.62 |
| $\Phi(\text{PSII})$ | $0.635 \pm 0.004$ a | 3.46 | $0.616 \pm 0.006$ a | 5.63 | $0.636 \pm 0.005$ a | 4.63 |
| NPQ | $0.291 \pm 0.007$ a | 12.42 | $0.275 \pm 0.010$ a | 19.53 | $0.287 \pm 0.008$ a | 15.60 |
| qP | $0.911 \pm 0.004$ a | 2.47 | $0.903 \pm 0.006$ a | 3.53 | $0.906 \pm 0.005$ a | 2.88 |
| $F_0$ | $0.384 \pm 0.005$ a | 6.52 | $0.391 \pm 0.003$ a | 4.59 | $0.382 \pm 0.005$ a | 6.47 |
| $F_m$ | $1.742 \pm 0.015$ a | 4.87 | $1.693 \pm 0.021$ a | 6.92 | $1.761 \pm 0.023$ a | 7.12 |

**Table 3.** Tomato leaf infrared gas analysis measurements on day 40 during growth under poly-chromatic spectrum. P_N_, net photosynthetic rate (P_gross_ – R_D_); g_s_, stomatal conductance; WUE, water use efficiency; E, transpiration rate. CV, coefficient of variance; SEM, standard error of the mean. The statistical significance of differences is indicated with letters (repeated measures one-way ANOVA, P<0.1). n = 10 for each experiment except one BLUE experiment with n = 5.

|  | **VIOLET (peak at 400 nm)** |  | **INDIGO (peak at 420 nm)** |  | **BLUE (peak at 450 nm)** |  |
| --- | --- | --- | --- | --- | --- | --- |
| | Mean ± SEM | CV (%) | Mean ± SEM | CV (%) | Mean ± SEM | CV (%) |
| $P_N$ | 6.55 ± 0.31 a | 26.0 | 6.16 ± 0.41 a | 36.7 | 8.92 ± 0.32 b | 18.19 |
| $g_s$ | 250.6 ± 19.8 a | 43.2 | 165.6 ± 24.2 a | 80.1 | 231.0 ± 29.4 a | 63.7 |
| WUE | 1.62 ± 0.13 a | 43.7 | 2.36 ± 0.23 a | 53.0 | 3.73 ± 0.53 a | 71.4 |
| E | 4.57 ± 0.28 a | 34.1 | 3.30 ± 0.36 a | 60.2 | 3.52 ± 0.42 a | 59.2 |

At harvest, after 120 days of growth, when two thirds of the fruit were ripe, the green vegetative tissue and all of the fruit was harvested and measured. Interestingly, for both fresh (FW) and dry weight (DW) of leaf tissue, we found that shifting the wavebands in the spectrum from BLUE (450 nm) to any of the shorter wavebands increased the biomass (Table 4). The BLUE-related leaf weight loss was not due to fewer leaves on the plants compared to light with VIOLET (400 nm) and INDIGO (420 nm), but a reduction in leaf area, including the specific leaf area (leaf area per unit weight), i.e., smaller and thicker leaves for the spectrum with the BLUE (450 nm) waveband (Figure 3A-C).

**Figure 3.**
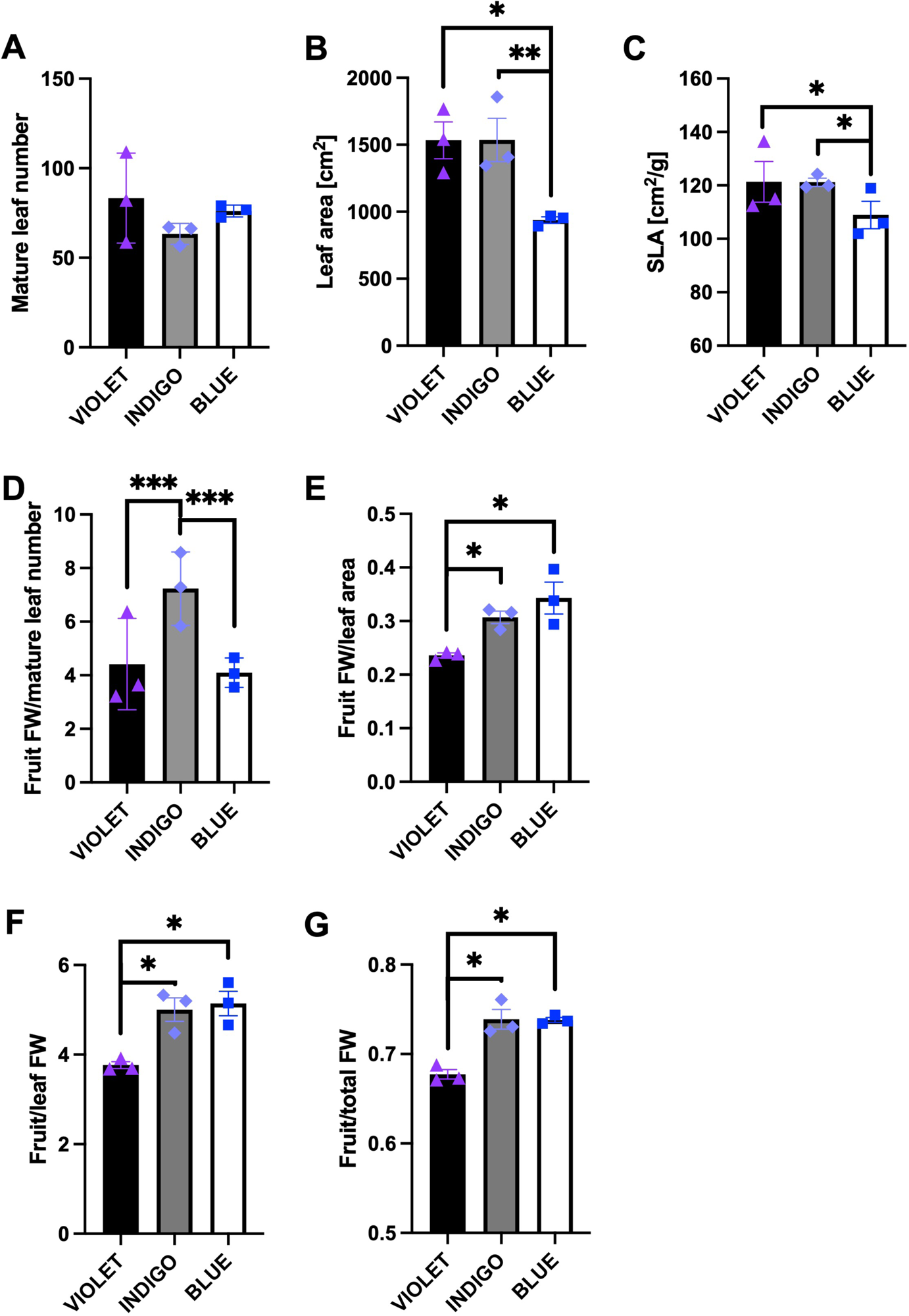
Tomato biomass measurements. Tomato leaf data and leaf/fruit ratio at harvest after growth under poly-chromatic conditions. Data points are the average values of each independent trial (3 trials per light treatment). (A) Mature leaf number. (B) Leaf area. (C) Specific leaf area, SLA. (D) Total fruit relative to mature leaf number. (E) Total fruit relative to leaf area. (F) Total fruit to leaf biomass ratio. (G) Total fruit to total biomass (all fruit plus stem and leaves) ratio. Error bars are S.E.M. The statistical significance of differences is indicated according to nested 1-way ANOVA analysis. *, *P* < 0.05; **, *P* < 0.01; ***, *P* < 0.001. n = 5 or 6 for each experiment.

**Table 4.**
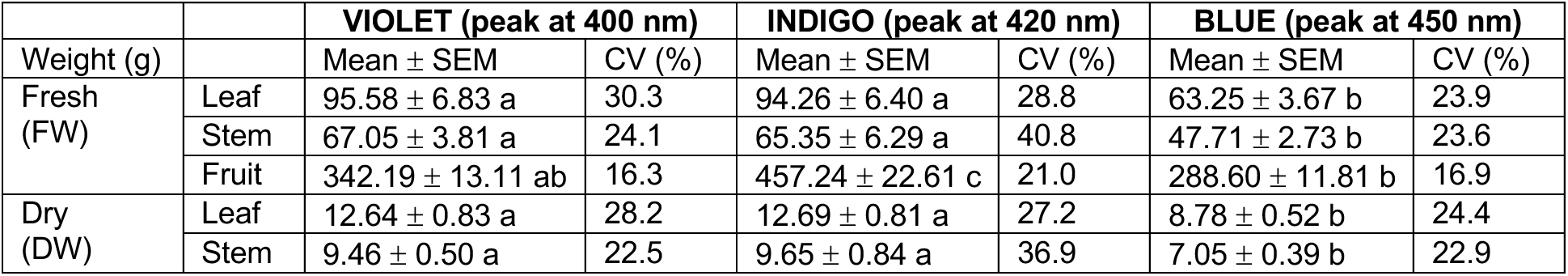
Mean biomass values of independent and triplicate experiments of tomato growth under poly-chromatic growth spectra with different blue wavebands. Fruit is total amount including unripe fruit. The statistical significance of differences is indicated with letters (repeated measures one-way ANOVA, P<0.05). n = 5 or 6 for each experiment.

The total fresh fruit yield for the spectrum with BLUE was significantly decreased compared to the spectrum with INDIGO, whereas VIOLET and INDIGO spectrum resulted in similar yield (Table 4). In contrast, however, the harvestable fruit yield and individual red fruit size was similar for all the light treatments (Table 5).

**Table 5.**
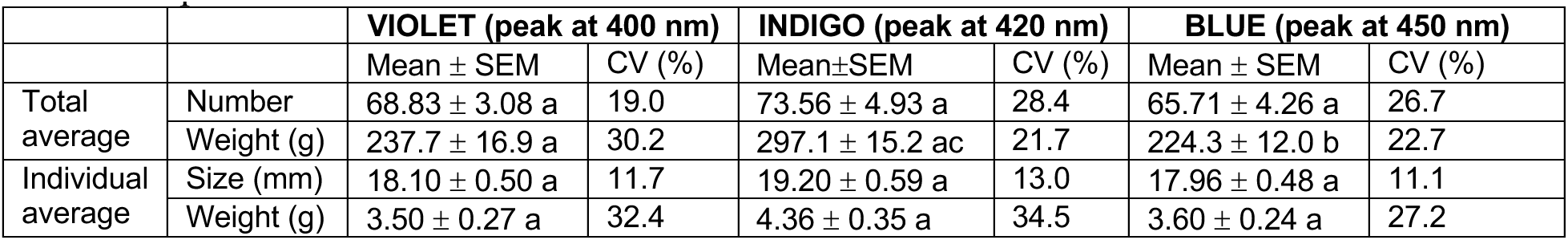
Number, size, and weight of harvestable fruit. Average biomass values for red fruit at harvest following growth under poly-chromatic light. Size is fruit diameter. The statistical significance of differences is indicated with letters (repeated measures one-way ANOVA, P<0.05). n = 5 or 6 for each experiment.

The lack of correlation between leaf biomass and fruit yield, prompted the calculation of fruit/leaf ratios. Both spectra with INDIGO and BLUE had higher total fruit/total FW and total fruit/leaf FW ratios than the spectrum with VIOLET, but only the spectrum with BLUE resulted in significantly higher fruit FW/leaf area ratio compared to VIOLET spectrum (Figure 3D-G). Thus, both INDIGO and BLUE spectrum had the most efficient fruit production with relation to green biomass whereas only the spectrum with BLUE was most efficient when measured against leaf area.

### 3.3. Leaf and fruit analysis

To analyze the response of plant pigment production to changes in blue spectral quality, both leaves and fruit were processed at harvest. We found no change in leaf chlorophyll and leaf total carotenoid content between the blue light treatments. The Chl a/b ratio however was significantly reduced for the light spectrum with BLUE (with a peak wavelength of 450 nm) compared to spectra with VIOLET (with a peak wavelength of 400 nm) and INDIGO (with a peak wavelength of 420 nm) (Table 6).

**Table 6.** Leaf pigment content. Tomato chlorophyll and carotenoid leaf pigment content measured at harvest (mg/mg FW) following growth under poly-chromatic conditions. TCC, total carotenoid content. The statistical significance of differences is indicated with letters (nested one-way ANOVA, P<0.05). n = 5 or 6 for each experiment.

| Pigment | VIOLET (peak at 400 nm) |  | INDIGO (peak at 420 nm) |  | BLUE (peak at 450 nm) |  |
| --- | --- | --- | --- | --- | --- | --- |
| | Mean ± SEM | CV (%) | Mean ± SEM | CV (%) | Mean ± SEM | CV (%) |
| Chl a | 1.96 ± 0.07 a | 14.4 | 1.98 ± 0.07 a | 14.7 | 1.79 ± 0.11 a | 24.8 |
| Chl b | 0.81 ± 0.04 a | 18.5 | 0.80 ± 0.03 a | 14.4 | 0.84 ± 0.05 a | 24.6 |
| Chl a/b | 2.48 ± 0.10 ab | 16.5 | 2.46 ± 0.04 a | 7.3 | 2.15 ± 0.06 c | 11.8 |
| TCC | 0.47 ± 0.02 a | 15.2 | 0.47 ± 0.02 a | 16.9 | 0.41 ± 0.03 a | 31.8 |

The pigment content and antioxidant activity of tomato fruits are important for tomato nutritional value. Between treatments, we found no difference in the content of the carotenoids beta-carotene and lycopene in red fruit. In contrast, there was a significantly elevated content of ASA in fruits exposed to the spectrum with BLUE (450 nm) relative to the spectra with VIOLET (400 nm) and INDIGO (420 nm). Finally, we found no difference in total phenolic acid (TPC) content and total soluble solids (TSS) between the blue light treatments (Table 7).

**Table 7.** Fruit physiological characterization. Pigment and metabolite content measurement of red fruit following growth under poly-chromatic spectrum. GAE, gallic acid equivalent; TPS, total phenol content; TSS, total soluble solids. The statistical significance of differences is indicated with letters (nested one-way ANOVA, P≤0.05). n = 5 or 6 for each experiment.

| Component | Unit | VIOLET (peak at 400 nm) |  | INDIGO (peak at 420 nm) |  | BLUE (peak at 450 nm) |  |
| --- | --- | --- | --- | --- | --- | --- | --- |
| | | Mean ± SEM | CV (%) | Mean ± SEM | CV (%) | Mean ± SEM | CV (%) |
| beta-Carotene | mg/g FW | 0.063 ± 0.005 a | 31.7 | 0.066 ± 0.004 a | 24.5 | 0.086 ± 0.005 a | 24.0 |
| Lycopene | | 0.591 ± 0.046 a | 30.9 | 0.778 ± 0.031 a | 15.6 | 0.834 ± 0.072 a | 34.6 |
| Ascorbic acid | | 28.31 ± 0.99 a | 25.7 | 22.62 ± 0.70 ab | 23.0 | 39.08 ± 1.57 c | 27.5 |
| TPC | mM GAE/g FW | 2.56 ± 0.07 a | 33.1 | 2.77 ± 0.06 a | 30.0 | 2.39 ± 0.06 a | 28.6 |
| TSS | °Brix | 3.22 ± 0.10 a | 13.5 | 3.61 ± 0.12 a | 13.7 | 3.72 ± 0.09 a | 10.4 |

## 4. Discussion

Blue light is essential for optimal plant growth and development, including photomorphogenesis, stomatal opening dynamics and chloroplast movement within cells or leaf tissue, which help to maximize photosynthesis. Blue light is high-energy light, relative to longer wavelengths, however, it is less efficiently used compared to red light in photosynthesis because it is immediately degraded to a lower energy level when absorbed by chlorophyll (Björn et al., 2009). In addition, blue light is absorbed by carotenoids and flavonoids that are pigments not directly involved in energy transmission (rather non-photochemical quenching). Nitride semiconductors are used to make short waveband (<540 nm) LEDs, and manufacturers typically focus on specific wavelengths that have sufficient commercial applications so that volume production can drive lower costs. Therefore, common LED product wavelengths include 365, 385, 405, 420 and 445 nm. LED physics and normal production variability leads to product peak wavelength variability of +/- 5 nm, a spectral full width at half maximum (FWHM) of approximately 20 nm, and a range of brightness bins (Dahl, 2010). Thus, the waveband variation in blue light LEDs offers the opportunity to optimize blue spectral quality for plant growth using sole-source lighting.

### 4.1. Photosynthetic light responses under mono- and polychromatic light

Blue light added to a red light background is known to enhance photosynthesis. To analyze the photosynthetic phenotype, tomato light response curves have often been generated following growth under different grow-light intensities, (e.g., Ke et al., 2023; Ke et al., 2022), changes in ambient CO_2_ content (Pan et al., 2020) or temperatures (Kong et al., 2021). Few reports were published on the tomato photosynthetic light response with relation to the light spectrum (and not including changes in blue spectral quality, e.g., Kaiser et al., 2019; Niu et al., 2023). Lanoue et al. (2017) compared monochromatic light response curves of the whole plant net exchange rate for blue (with a peak wavelength of 440 nm), green and red light in ‘Bonny Best’ tomato. Blue light resulted in the highest LCP and lowest net carbon exchange rate compared to green light and, in one study, also to red light (Lanoue et al., 2017; Lanoue et al., 2018). The reported curve values, e.g., LCP, were different from our findings, a discrepancy that could be due to cultivar difference or single leaf vs. whole plant measurement (Lanoue et al., 2018). We showed that blue wavebands are largely interchangeable for the estimated rate of photochemical efficiency of ‘Micro Tom’ under mono- and polychromatic conditions (Figure 2C-D). This is interesting since Sager et al. (1988) measured a low relative quantum efficiency of 400 nm compared to 420 nm and 450 nm, however the data were obtained for isolated phytochrome *in vitro* (Sager et al., 1988). More cultivars should be tested in order to conclude with finality if the blue spectrum is crucial to the photosynthetic phenotype of tomato.

For poly-chromatic spectrum with BLUE (peak at 450 nm), we found that the net photosynthetic rate during growth was inversely correlated with final biomass (Figure 3 and Table 4). This correlation has also been observed in many other blue or UV-A light studies. There are several photoreceptors responsible for light perception in the 380-450 nm spectrum (e.g., cryptochrome, phototropin, phytochrome A). Short wavelength light signaling is not a direct signaling pathway at the molecular level and varied responses to light supplemented by UV-A or blue light are therefore observed in different crop species (Verdaguer et al., 2017). The relatively high P_N_ for BLUE during growth was in contrast to the corresponding lower P_N_ value for the light response curve, likely explained by younger age of plants cultivated under FL prior to light response analysis (vs. growth trials under LED lights).

### 4.2. Blue spectral quality affects plant size

In tomato crop development, spectral quality was often studied with regards to the relative proportions of blue, green, red and far-red light. Comparison of specific wavebands within the shorter-wavelength blue (∼400-450 nm) spectrum has rarely been reported on. Blue supplemental light of longer wavelength blue, 455 nm and 470 nm, were compared for tomato grown under HPS light in the greenhouse and the effects (height, pigment content, leaf area) were shown to be cultivar dependent (Brazaitytė and Kasiulevičiūtė, 2013). In another study, the comparison of poly-chromatic spectra with/without 380 nm blue light revealed that 380 nm had a positive effect on shoot dry weight and leaf area of tomato (Brazaitytė et al., 2010). Finally, when blue (450 nm) was replaced with UV-A (370 nm) light in an RB LED fixture, both leaf area and plant height was increased, not SLA, and chlorophyll and phenolic content in leaves was reduced, suggesting growth optimization via morphological, not photosynthetic, adjustment (the fruit yield was not analyzed) (Zhang et al., 2020).

We found that the blue spectral quality evaluated affected the leaf area, which was significantly smaller for poly-chromatic spectrum with BLUE (450 nm) compared to VIOLET (400 nm) and INDIGO (420 nm). There are conflicting data in the literature whether leaf area is influenced by blue light and the blue/red ratio in tomato. Adding blue to a red light background increased leaf number and leaf area in several tomato cultivars and species (Hernández et al., 2016; Ouzounis et al., 2016), whereas another analysis (of young plants) showed that only dry weight partitioning to the leaf, compared to stem, was increased (Kalaitzoglou et al., 2021). For ‘Micro Tom’ in particular, in one case, a higher blue proportion resulted in increased SLA, not leaf number and area (Ke et al., 2022), in another case, a low leaf number and small leaf area (Vitale et al., 2022) – a discrepancy likely caused by different light recipes used (specifically, the quantities of green and far-red light). The variation in phenotypes observed are likely due to the large natural variation in tomato. During tomato domestication, thousands of cultivars were selected for, all adapted to specific environments, resulting in a rich diversity of phenotypes related to plant-environment interactions (Tripodi, 2022). In Arabidopsis, natural variation was demonstrated for blue light-associated anthocyanin content in leaves linked to biomass accumulation (Yavari et al., 2021). In tomato, blue (450 nm) fluence rate correlated negatively with plant height and SLA and there was a threshold (15-20% PFD blue) for positive correlation with biomass (Utasi et al., 2023). Leaf pigment-dependent biomass accumulation is not well studied in tomato but direct evidence for the role of blue light perception in tomato leaf size was shown for the tomato cryptochrome mutant that had a larger leaf area compared to the wild type (Liu et al., 2018). Our finding that the shorter wavelengths of blue light resulted in larger leaf areas, is in agreement with the fact that UV-A (i.e., short wavelength blue) radiation promotes tomato plant growth through morphological adaptation (increase in light interception) (Zhang et al., 2020).

### 4.3. Biomass yield and fruit yield efficiency

Despite the common result of increased plant compactness following high blue light dosage, species-specific biomass outcomes are known for blue light treatments (Huché-Thélier et al., 2016). In agreement with the leaf size data, we found that the biomass for the BLUE (450 nm) condition was significantly less than for VIOLET (400 nm) and INDIGO (420 nm) light. This is consistent with an earlier study that showed that the addition of short wavelength (380 nm) light to a white spectrum increased the dry-weight of young tomato plants (Brazaitytė et al., 2010). We also found that INDIGO resulted in the highest total fruit yield compared to VIOLET and BLUE, and BLUE had the lowest total fruit yield, and these results were similar to the mass of harvestable fruit (Table 4 and 6).

When fruit yield efficiency (harvest index) was calculated in relation to total biomass or total leaf mass or area, both INDIGO (420 nm) and BLUE (450 nm) had higher yield efficiency than VIOLET (400 nm). INDIGO, however, was the only condition where the yield efficiency relative to leaf number was significantly elevated, indicating a specific influence of INDIGO on leaf development.

Blue spectral diversity, similar to what we tested, was previously applied to growth of kale, but no changes in biomass were found, only for leaf pigment composition (Ashenafi et al., 2023). For another leafy crop, perilla, blue light with peak wavelengths of 385 nm, 415 nm and 430 nm, was supplemented to white LED light and the photosynthetic rate, leaf area and dry weight was increased but only in the red cultivar, not the green. For both cultivars, the biomass correlated with the photosynthetic rate and also biochemical changes (Nguyen and Oh, 2022). Finally, varied responses in biomass and photosynthetic rate were found for three unrelated leafy greens supplemented with peak wavelengths of either 450 nm or 420 nm plus 440 nm blue light (Taulavuori et al., 2018). Thus, efforts will have to be continued in order to catalogue and understand blue spectral specific effects on crop growth and quality in indoor growth operations.

### 4.4. Blue spectral quality is reflected in the Chl a/b ratio

The Chl *a*/*b* ratio reflects fluctuations in the light environment and is a species specific response (Murchie and Horton, 1998; Schöttler and Tóth, 2014). In several species, including tomato, blue light and high fluence rate can promote “sun-type” leaves and elevate Chl *a*/*b* (Ballottari et al., 2007; Fang et al., 2021; Hogewoning et al., 2007; Wang et al., 2015; Wang et al., 2016; Wei et al., 2017). Since Chl *b* is primarily bound by the light harvesting complexes of PSII (LHCII), the Chl *a*/*b* ratio correlates with the stoichiometry of the photosystems, the PSII/PSI ratio, and alterations in Chl *a*/*b* is therefore evidence of active light acclimation (Kim et al., 1993; Holtzegel, 2016; Kouřil et al., 2013; Pfannschmidt et al., 1999). Chl *a*/*b* ratios have often been compared following changes in light irradiance or the blue/red light ratio, including in ‘Micro Tom’ tomato, where a higher proportion of blue light lowered Chl *a*/*b* (Vitale et al., 2022). Our result that BLUE (450 nm) lowered Chl *a*/*b* compared to VIOLET (400 nm) and INDIGO (420 nm) in a poly-chromatic spectrum indicates even subtle shifting of blue wavebands are sensed by the plant and photosynthesis is adjusted accordingly (it is a sensitive and dynamic system). We did not observe other changes in pigment content of the leaves, in contrast to Vitale et al. (2022), which could be due to later leaf harvesting compared to Vitale et al. (2022).

### 4.5. 450 nm blue light maximizes ASA content

An interesting significant effect of long wavelength blue light was the higher ASA content under the BLUE (450 nm) treated plants compared to VIOLET (400 nm) and INDIGO (420 nm). This result agrees with the recent identification of a new blue light photoreceptor in tomato, named PLP for PAS/LOV protein. Upon blue light perception, PLP is bound to GDP-L-galactose-phosphorylase and ASA biosynthesis is de-repressed (Bournonville et al., 2023). The LOV domain is a well-known chromophore in blue light photoreceptors (e.g., phototropins) and the LOV absorbance spectrum peaks at 450 nm (Christie et al., 1999). When blue light of 470 nm was compared to red light, there was no change in tomato ASA content (Xiao et al., 2022). Taking these findings into context with the PLP molecular mechanism, likely only blue light with a peak of 450 nm can fully de-activate PLP and promote ASA biosynthesis, compared to other blue wavelengths. The quantity of blue light is also of importance since a higher proportion of blue promoted ASA content (21% vs. 33% PFD blue light) in ‘Micro Tom’ tomato (Vitale et al., 2022). Finally, blue light (with a peak wavelength of 450 nm) application following tomato fruit harvest also resulted in increased ASA content (Ntagkas et al., 2019).

### 4.6. Pigment content and nutritional value

Compared to greenhouse cultivation, i.e., using sunlight, the manipulation of the light spectrum using sole-source lighting has great potential for modulation of tomato phytochemical content including pigments (Dzakovich et al., 2017; Holopainen et al., 2018). Greenhouse data are variable, e.g., earlier studies of spectral influence on secondary metabolites of ‘Micro Tom’ tomato fruits showed that both blue (with a peak wavelength of 430 nm) and red (with a peak wavelength of 660 nm) supplemental light elevated the content of lycopene, lutein and beta-carotene, but blue light had the strongest effect (Xie et al., 2019). In another study, when blue light (with a peak wavelength of 450 nm) was applied as monochromatic illumination the tomatoes had reduced lycopene content (Li et al., 2021). Using UV (with a peak of 352 nm) supplemented fluorescent lighting, the nutritional quality of two dwarf tomato varieties was altered for multiple parameters including ASA (Kobayashi and Tabuchi, 2022). Finally, it was shown for red-fruited F1 hybrid cultivars that lamp types with the highest blue light percentage increased lycopene content (Alsina et al., 2022). For ‘Micro Tom’, Vitale et al. (2022) found that TPC was inversely correlated with the amount of blue light. We did not observe any changes in tomato TPC, indicating that changes in TPC only occur following manipulation of blue light intensity but not spectral quality.

### 4.7. Conclusions

In conclusion, our work revealed that blue spectral quality affects the growth and development of the dwarf tomato cultivar ‘Micro Tom’. While the photochemical efficiency was consistent across the examined wavebands, the net photosynthetic rate varied with blue spectral changes. This suggests that factors beyond photochemical efficiency play a role in determining the overall photosynthetic performance of tomato. The results also indicated distinct advantages associated with VIOLET (peak at 400 nm) and INDIGO (peak at 420 nm) light. Plants exposed to these colors exhibited larger leaf area and biomass, highlighting a potential for enhanced growth and productivity. The harvest index, a key parameter indicating the efficiency of resource allocation, was highest for spectra with INDIGO (peak at 420 nm) and BLUE (peak at 450 nm). This underscored the effectiveness of blue wavebands in promoting the development of reproductive structures (sink tissue) relative to foliage. The BLUE (450 nm) light condition also resulted in a lower Chl *a*/*b* ratio, indicating a specific role in adaptation to light absorption and photochemical quenching. Moreover, the higher ASA content observed for BLUE indicated a role in the biosynthesis of secondary metabolites. No other nutritional alterations were detected in the tomato fruits, suggesting a distinct role for blue spectral quality in tomato fruit nutritional composition. Thus, our study revealed the interplay between blue spectral quality and plant physiology, demonstrating the nuanced effects that blue light wavelengths can exert on growth, development, and biochemical responses. Our findings underscore the need for a holistic understanding of light-plant interactions in horticultural practices to maximize both yield and nutritional quality in indoor crop production.

## Author Contributions

Experimental design: E.K. Experimental work: M.S.G. Data analysis: M.S.G. and E.K. Writing: E.K. All authors contributed to the article and approved the submitted version.

## Acknowledgment

We are grateful for the technical assistance from Richard R. Neal. We thank A.J. Both and Neil S. Mattson for helpful discussions.

## References

Ainsworth, E.A. and K.M. Gillespie. 2007. Estimation of total phenolic content and other oxidation substrates in plant tissues using Folin–Ciocalteu reagent. Nature Protocols, 2: 875. doi: 10.1038/nprot.2007.102

Alsina, I., I. Erdberga, M. Duma, R. Alksnis and L. Dubova. 2022. Changes in greenhouse grown tomatoes metabolite content depending on supplemental light quality. Front. Nutr., 9: 830186. doi: 10.3389/fnut.2022.830186

AOAC. 2006. AOAC official method 967.21 – ascorbic acid in vitamin preparations and juices: 2,6-dichloroindophenol titrimetric method. Official methods of analysis of the Association of Official Analytical Chemists, 19–20.

Artés-Hernández, F., N. Castillejo and L. Martínez-Zamora. 2022. UV and visible spectrum LED lighting as abiotic elicitors of bioactive compounds in sprouts, microgreens, and baby leaves - a comprehensive review including their mode of action. Foods, 11: 265. doi: 10.3390/foods11030265

Ashenafi, E.L., M.C. Nyman, J.M. Holley and N.S. Mattson. 2023. The influence of LEDs with different blue peak emission wavelengths on the biomass, morphology, and nutrient content of kale cultivars. Sci. Hortic., 317: 111992. doi: 10.1016/j.scienta.2023.111992

Ballottari, M., L. Dall’osto, T. Morosinotto and R. Bassi. 2007. Contrasting behavior of higher plant photosystem I and II antenna systems during acclimation. J. Biol. Chem., 282: 8947– 8958. doi: 10.1074/jbc.M606417200

Björn, L.O., G.C. Papageorgiou, R.E. Blankenship and Govindjee. 2009. A viewpoint: Why chlorophyll a. Photosynth. Res., 99: 85–98. doi: 10.1007/s11120-008-9395-x

Bournonville, C. et al. 2023. Blue light promotes ascorbate synthesis by deactivating the PAS/LOV photoreceptor that inhibits GDP-l-galactose phosphorylase. Plant Cell, 2615–2634. doi: 10.1093/plcell/koad108

Brazaitytė, A. et al. 2010. The effect of light-emitting diodes lighting on the growth of tomato transplants. Zemdirbyste-Agriculture, 97: 89–98.

Brazaitytė, A. and A. Kasiulevičiūtė. 2013. The effects of HPS lamp supplementation with blue light-emitting diodes on the growth of two tomato hybrid transplants. Rural Dev., 6: 49–53.

Christie, J.M., M. Salomon, K. Nozue, M. Wada and W.R. Briggs. 1999. LOV (light, oxygen, or voltage) domains of the blue-light photoreceptor phototropin (nph1): Binding sites for the chromophore flavin mononucleotide. Proc. Natl. Acad. Sci. USA, 96: 8779–8783. doi: 10.1073/pnas.96.15.8779

Cosgrove, D.J. 1981. Rapid suppression of growth by blue light: Occurrence, time course, and general characteristics. Plant Physiol., 67: 584–590. doi: 10.1104/pp.67.3.584

Dahl, R. 2010. Light emitting diodes – choosing the right LED for the job. Tech Briefs, www.techbriefs.com. SAE Media Group

De Carbonnel, M., J.M. Stormonth-Darling, W. Liu, D. Kuziak and M.A. Jones. 2022. Realising the environmental potential of vertical farming systems through advances in plant photobiology. Biology (Basel), 11: 922. doi: 10.3390/biology11060922

Dzakovich, M.P., C. Gómez, M.G. Ferruzzi and C.A. Mitchell. 2017. Chemical and sensory properties of greenhouse tomatoes remain unchanged in response to red, blue, and far red supplemental light from light-emitting diodes. HortSci., 52: 1734–1741. doi: 10.21273/hortsci12469-17

Fang, L. et al. 2021. Plant growth and photosynthetic characteristics of soybean seedlings under different LED lighting quality conditions. *J*. Plant Growth Regulation, 40: 668–678. doi: 10.1007/s00344-020-10131-2

FAOSTAT. 2021. Food and agriculture statistics. Food and agriculture organization of the united nations. Available online: https://www.fao.org/faostat/en/#data/QCL (accessed on 26 September 2023).

Fenech, M., I. Amaya, V. Valpuesta and M.A. Botella. 2018. Vitamin C content in fruits: Biosynthesis and regulation. Front. Plant Sci., 9: 2006. doi: 10.3389/fpls.2018.02006

He, R. et al. 2022. Supplemental blue light frequencies improve ripening and nutritional qualities of tomato fruits. Front. Plant Sci., 13: 888976. doi: 10.3389/fpls.2022.888976

Hernández, R., T. Eguchi, M. Deveci and C. Kubota. 2016. Tomato seedling physiological responses under different percentages of blue and red photon flux ratios using LEDs and cool white fluorescent lamps. Sci. Hortic., 213: 270–280. doi: 10.1016/j.scienta.2016.11.005

Hikosaka, S. (2018) Production of value-added plants. In: Smart Plant Factory, Springer Singapore, Singapore, pp. 325–351.

Hoagland, D.R., and D.I. Arnon. 1950. The water-culture method for growing plants without soil. Circular. California agricultural experiment station, 347 (2nd edit).

Hogewoning, S.W. et al. 2007. Plant physiological acclimation to irradiation by light-emitting diodes (LEDs). Acta Hortic., 183–191. doi: 10.17660/actahortic.2007.761.23

Hogewoning, S.W., G. Trouwborst, H. Maljaars, H. Poorter, W. Van Ieperen and J. Harbinson. 2010. Blue light dose-responses of leaf photosynthesis, morphology, and chemical composition of *Cucumis sativus* grown under different combinations of red and blue light. J. Expt. Bot., 61: 3107–3117. doi: 10.1093/jxb/erq132

Holopainen, J.K., M. Kivimäenpää and R. Julkunen-Tiitto. 2018. New light for phytochemicals. Trends Biotechnol., 36: 7–10. doi: 10.1016/j.tibtech.2017.08.009

Holtzegel, U. 2016. The LHC family of *Arabidopsis thaliana*. Endocytobiosis Cell Res, 27: 71– 89.

Huché-Thélier, L., L. Crespel, J.L. Gourrierec, P. Morel, S. Sakr and N. Leduc. 2016. Light signaling and plant responses to blue and UV radiations—perspectives for applications in horticulture. Environ. Expt. Bot., 121: 22–38. doi: 10.1016/j.envexpbot.2015.06.009

Imada, K., S. Tanaka, Y. Ibaraki, K. Yoshimura and S. Ito. 2014. Antifungal effect of 405-nm light on *Botrytis cinerea*. Lett. Appl. Microbiol., 59: 670–676. doi: 10.1111/lam.12330

Kaiser, E., T. Ouzounis, H. Giday, R. Schipper, E. Heuvelink and L.F.M. Marcelis. 2018. Adding blue to red supplemental light increases biomass and yield of greenhouse-grown tomatoes, but only to an optimum. Front. Plant Sci., 9: 2002. doi: 10.3389/fpls.2018.02002

Kaiser, E., K. Weerheim, R. Schipper and J.A. Dieleman. 2019. Partial replacement of red and blue by green light increases biomass and yield in tomato. Sci. Hortic., 249: 271–279. doi: 10.1016/j.scienta.2019.02.005

Kalaitzoglou, P. et al. 2021. Unraveling the effects of blue light in an artificial solar background light on growth of tomato plants. Environ. Expt. Bot., 184: 104377. doi: 10.1016/j.envexpbot.2021.104377

Karlicek, R.F. 2019. New research LED fixture for plant growth. GLASE Technical Article, www.glase.org.

Ke, X., H. Yoshida, S. Hikosaka and E. Goto. 2022. Optimization of photosynthetic photon flux density and light quality for increasing radiation-use efficiency in dwarf tomato under LED light at the vegetative growth stage. Plants, 11: 121. doi: 10.3390/plants11010121

Ke, X., H. Yoshida, S. Hikosaka and E. Goto. 2023. Photosynthetic photon flux density affects fruit biomass radiation-use efficiency of dwarf tomatoes under LED light at the reproductive growth stage. Front. Plant Sci., 14: doi: 10.3389/fpls.2023.1076423

Kim, J.H., R.E. Glick and A. Melis. 1993. Dynamics of photosystem stoichiometry adjustment by light quality in chloroplasts. Plant Physiol., 102: 181–190. doi: 10.1104/pp.102.1.181

Kobayashi, T. and T. Tabuchi. 2022. Tomato cultivation in a plant factory with artificial light: Effect of uv-a irradiation during the growing period on yield and quality of ripening fruit. Horticulture J., 91: 16–23. doi: 10.2503/hortj.UTD-272

Kolmos, E., R.R. Neal, A. Tuzikas and R.F. Karlicek. 2021. Infra-red gas analyzer with custom LED module attachment for multi-spectral photosynthetic analyses. GLASE Technical Article, www.glase.org

Kong, L., Y. Wen, X. Jiao, X. Liu and Z. Xu. 2021. Interactive regulation of light quality and temperature on cherry tomato growth and photosynthesis. Environ. Expt. Bot., 182: 104326. doi: 10.1016/j.envexpbot.2020.104326

Kouřil, R., E. Wientjes, J.B. Bultema, R. Croce and E.J. Boekema. 2013. High-light vs. low-light: Effect of light acclimation on photosystem II composition and organization in *Arabidopsis thaliana*. Biochim. Biophys. Acta, 1827: 411–419. doi: 10.1016/j.bbabio.2012.12.003

Lanoue, J., E.D. Leonardos, S. Khosla, X. Hao and B. Grodzinski. 2018. Effect of elevated CO_2_ and spectral quality on whole plant gas exchange patterns in tomatoes. PLoS One, 13: e0205861. doi: 10.1371/journal.pone.0205861

Lanoue, J., E.D. Leonardos, X. Ma and B. Grodzinski. 2017. The effect of spectral quality on daily patterns of gas exchange, biomass gain, and water-use-efficiency in tomatoes and Lisianthus: An assessment of whole plant measurements. Front Plant Sci, 8: 1076. doi: 10.3389/fpls.2017.01076

Li, Y., C. Liu, Q. Shi, F. Yang and M. Wei. 2021. Mixed red and blue light promotes ripening and improves quality of tomato fruit by influencing melatonin content. Environ. Expt. Bot., 185: 104407. doi: 10.1016/j.envexpbot.2021.104407

Lichtenthaler, H.K. and C. Buschmann. 2001. Chlorophylls and carotenoids: Measurement and characterization by UV-VIS spectroscopy. Curr. Protocols Food Anal. Chem., 1: F4.3.1–F4.3.8. doi: 10.1002/0471142913.faf0403s01

Liu, C.C. et al. 2018. Tomato CRY1a plays a critical role in the regulation of phytohormone homeostasis, plant development, and carotenoid metabolism in fruits. Plant Cell Environ., 41: 354–366. doi: 10.1111/pce.13092

Lobo, F.D.A. et al. 2013. Fitting net photosynthetic light-response curves with Microsoft Excel—a critical look at the models. Photosynthetica, 51: 445–456. doi: 10.1007/s11099-013-0045-y

Long, S.P., X.G. Zhu, S.L. Naidu and D.R. Ort. 2006. Can improvement in photosynthesis increase crop yields. Plant Cell Environ., 29: 315–330. doi: 10.1111/j.1365-3040.2005.01493.x

Matsuda, R., K. Ohashi-Kaneko, K. Fujiwara, E. Goto and K. Kurata. 2004. Photosynthetic characteristics of rice leaves grown under red light with or without supplemental blue light. Plant Cell Physiol., 45: 1870–1874. doi: 10.1093/pcp/pch203

Mitchell, C.A. 2022. History of controlled environment horticulture: Indoor farming and its key technologies. HortSci., 57: 247–256. doi: 10.21273/hortsci16159-21

Mochizuki, T., Y. Onda, E. Fujiwara, M. Wada and Y. Toyoshima. 2004. Two independent light signals cooperate in the activation of the plastid psbD blue light-responsive promoter in Arabidopsis. FEBS Lett., 571: 26–30. doi: 10.1016/j.febslet.2004.06.052

Muneer, S., E.J. Kim, J.S. Park and J.H. Lee. 2014. Influence of green, red and blue light emitting diodes on multiprotein complex proteins and photosynthetic activity under different light intensities in lettuce leaves (*Lactuca sativa* L.). Int. J. Mol. Sci., 15: 4657–4670. doi: 10.3390/ijms15034657

Murchie, E.H. and P. Horton. 1998. Contrasting patterns of photosynthetic acclimation to the light environment are dependent on the differential expression of the responses to altered irradiance and spectral quality. Plant Cell Environ., 21: 139–148. doi: 10.1046/j.1365-3040.1998.00262.x

Nagata, M. and I. Yamashita. 1992. Simple method for simultaneous determination of chlorophyll and carotenoids in tomato fruit. Nippon Shokuhin Kogyo Gakkaishi, 39: 925– 928. doi: 10.3136/nskkk1962.39.925

Nguyen, L.T.K. and M.M. Oh. 2022. Growth and biochemical responses of green and red Perilla supplementally subjected to UV-A and deep-blue LED lights. Photochem. Photobiol., 98: 1332–1342. doi: 10.1111/php.13614

Niu, Y., H. Lyu, X. Liu, M. Zhang and H. Li. 2023. Photosynthesis prediction and light spectra optimization of greenhouse tomato based on response of red–blue ratio. Sci. Hortic., 318: 112065. doi: 10.1016/j.scienta.2023.112065

Nozue, H. and Gomi, M. (2018) Usefulness of broad-spectrum white LEDs to envision future plant factory. In: Smart Plant Factory, Springer Singapore, Singapore, pp. 197–210.

Ntagkas, N., R.C.H. De Vos, E.J. Woltering, C.C.S. Nicole, C. Labrie and L.F.M. Marcelis. 2020. Modulation of the tomato fruit metabolome by LED light. Metabolites, 10: E266. doi: 10.3390/metabo10060266

Ntagkas, N., E. Woltering, C. Nicole, C. Labrie and L.F.M. Marcelis. 2019. Light regulation of vitamin C in tomato fruit is mediated through photosynthesis. Environ. Expt. Botany, 158: 180–188. doi: 10.1016/j.envexpbot.2018.12.002

Oh, S. and C. Lu. 2022. Vertical farming-smart urban agriculture for enhancing resilience and sustainability in food security. J. Hort. Sci. Biotech., 1–8. doi: 10.1080/14620316.2022.2141666

Ouzounis, T., E. Heuvelink, Y. Ji, H.J. Schouten, R.G.F. Visser and L.F.M. Marcelis. 2016. Blue and red LED lighting effects on plant biomass, stomatal conductance, and metabolite content in nine tomato genotypes. Acta Hort., 251–258. doi: 10.17660/actahortic.2016.1134.34

Ouzounis, T., E. Rosenqvist and C.-O. Ottosen. 2015. Spectral effects of artificial light on plant physiology and secondary metabolism: A review. HortSci., 50: 1128–1135. doi: 10.21273/hortsci.50.8.1128

Pan, T. et al. 2020. Increased CO_2_ and light intensity regulate growth and leaf gas exchange in tomato. Physiol. Plant, 168: 694–708. doi: 10.1111/ppl.13015

Paradiso, R. and S. Proietti. 2021. Light-quality manipulation to control plant growth and photomorphogenesis in greenhouse horticulture: The state of the art and the opportunities of modern LED systems. *J*. Plant Growth Regulation, doi: 10.1007/s00344-021-10337-y

Pfannschmidt, T., A. Nilsson and J.F. Allen. 1999. Photosynthetic control of chloroplast gene expression. Nature, 397: 625–628. doi: 10.1038/17624

Retana-Cordero, M., S. Humphrey and C. Gómez. 2022. Effect of radiation quality and relative humidity on intumescence injury and growth of tomato seedlings. HortSci., 57: 1257–1266. doi: 10.21273/hortsci16712-22

Sager, J.C., W.O. Smith, J.L. Edwards and K.L. Cyr. 1988. Photosynthetic efficiency and phytochrome photoequilibria determination using spectral data. Transactions ASAE, 31: 1882–1889.

Salehi, B. et al. 2019. Beneficial effects and potential risks of tomato consumption for human health: An overview. Nutrition, 62: 201–208. doi: 10.1016/j.nut.2019.01.012

Schöttler, M.A. and S.Z. Tóth. 2014. Photosynthetic complex stoichiometry dynamics in higher plants: Environmental acclimation and photosynthetic flux control. Front. Plant Sci., 5: 188. doi: 10.3389/fpls.2014.00188

Scott, J.W. 1989. Micro-Tom - a miniature dwarf tomato. FL Agric. Expt. Sta. Circ., 370: 1–6.

Sharathkumar, M., E. Heuvelink and L.F.M. Marcelis. 2020. Vertical farming: Moving from genetic to environmental modification. Trends Plant Sci., doi: 10.1016/j.tplants.2020.05.012

Sytar, O., Zivcak, M., Brestic, M., Toutounchi, P.M. and Allakhverdiev, S.I. (2021) Plasticity of the photosynthetic energy conversion and accumulation of metabolites in plants in response to light quality. In: Photosynthesis: Molecular approaches to solar energy conversion: Advances in photosynthesis and respiration, Springer International Publishing, Cham, pp. 533–563.

Taulavuori, K., A. Pyysalo, E. Taulavuori and R. Julkunen-Tiitto. 2018. Responses of phenolic acid and flavonoid synthesis to blue and blue-violet light depends on plant species. Environ. Expt. Bot., 150: 183–187. doi: 10.1016/j.envexpbot.2018.03.016

Tripodi, P. 2022. Next generation sequencing technologies to explore the diversity of germplasm resources: Achievements and trends in tomato. Comput. Struct. Biotechnol. J., 20: 6250– 6258. doi: 10.1016/j.csbj.2022.11.028

Utasi, L., V. Kovács, Z. Gulyás, T. Marcek, T. Janda and E. Darko. 2023. Threshold or not: Spectral composition and light-intensity dependence of growth and metabolism in tomato seedlings. Sci. Hortic., 313: 111946. doi: 10.1016/j.scienta.2023.111946

Verdaguer, D., M.A. Jansen, L. Llorens, L.O. Morales and S. Neugart. 2017. UV-A radiation effects on higher plants: Exploring the known unknown. Plant Sci., 255: 72–81. doi: 10.1016/j.plantsci.2016.11.014

Vitale, E., V. Velikova, T. Tsonev, G. Costanzo, R. Paradiso and C. Arena. 2022. Manipulation of light quality is an effective tool to regulate photosynthetic capacity and fruit antioxidant properties of *Solanum lycopersicum* L. cv. ‘Microtom’ in a controlled environment. PeerJ., 10: e13677. doi: 10.7717/peerj.13677

Wang, J., W. Lu, Y. Tong and Q. Yang. 2016. Leaf morphology, photosynthetic performance, chlorophyll fluorescence, stomatal development of lettuce (*Lactuca sativa* L.) exposed to different ratios of red light to blue light. Front. Plant Sci., 7: 250. doi: 10.3389/fpls.2016.00250

Wang, S. et al. 2022. Response of tomato fruit quality depends on period of LED supplementary light. Front. Nutr., 9: 833723. doi: 10.3389/fnut.2022.833723

Wang, X.Y., X.M. Xu and J. Cui. 2015. The importance of blue light for leaf area expansion, development of photosynthetic apparatus, and chloroplast ultrastructure of *Cucumis sativus* grown under weak light. Photosynthetica, 53: 213–222. doi: 10.1007/s11099-015-0083-8

Wei, H., W. Xiaoxiao, P. Min, L. Xiaoying, G. Lijun and X. Zhigang. 2017. Effect different spectral LED on photosynthesis and distribution of photosynthate of cherry tomato seedlings. 2017 14th China International Forum on Solid State Lighting: International Forum on Wide Bandgap Semiconductors China, doi: 10.1109/ifws.2017.8245979

Xiao, L., T. Shibuya, T. Watanabe, K. Kato and Y. Kanayama. 2022. Effect of light quality on metabolomic, ionomic, and transcriptomic profiles in tomato fruit. Int. J. Mol. Sci., 23: 13288. doi: 10.3390/ijms232113288

Xie, B.-X. et al. 2019. Supplemental blue and red light promote lycopene synthesis in tomato fruits. J. Integrative Agric., 18: 590–598. doi: 10.1016/s2095-3119(18)62062-3

Yavari, N., R. Tripathi, B.S. Wu, S. Macpherson, J. Singh and M. Lefsrud. 2021. The effect of light quality on plant physiology, photosynthetic, and stress response in *Arabidopsis thaliana* leaves. PLoS One, 16: e0247380. doi: 10.1371/journal.pone.0247380

Zelkind, M., Livingston, T. and Verlage, V. (2022) Indoor production of tomatoes. In: Plant Factory Basics, Applications and Advances, Elsevier, pp. 295–305.

Zhang, Y. et al. 2020. UVA radiation promotes tomato growth through morphological adaptation leading to increased light interception. Environ. Expt. Bot., 176: 104073. doi: 10.1016/j.envexpbot.2020.104073

Zhen, S., M. Van Iersel and B. Bugbee. 2021. Why far-red photons should be included in the definition of photosynthetic photons and the measurement of horticultural fixture efficacy. Front. Plant Sci., 12: 693445. doi: 10.3389/fpls.2021.693445

